# Microclimate is a strong predictor of the native and invasive plant-associated soil microbiota on San Cristóbal Island, Galápagos archipelago

**DOI:** 10.1101/2022.04.05.487164

**Authors:** Alexi A. Schoenborn, Sarah M. Yannarell, Caroline T. MacVicar, Noelia N. Barriga-Medina, Meng Markillie, Hugh Mitchell, Kevin S. Bonham, Antonio Leon-Reyes, Diego Riveros-Iregui, Vanja Klepac-Ceraj, Elizabeth A. Shank

## Abstract

Understanding the major drivers that influence soil bacterial and fungal communities is essential to mitigate the impacts of human activity on vulnerable ecosystems, like those found on the Galápagos Islands. Located ~1000 km off the coast of Ecuador, the volcanically formed islands are situated within distinct oceanic currents, which provide seasonal weather patterns and unique microclimates within small spatial scales across the islands. Although much is known about the impacts of human activity, such as climate change and invasive plant species, on above ground biodiversity of the Galápagos Islands, little is known about the resident soil microbial communities and the drivers that shape these communities. Here, our goal was to investigate the bacterial and fungal communities found in soil located in three distinct microclimates: Mirador (arid), Cerro Alto (transition zone), and El Junco (humid), and associated with native and invasive plant types. At each site, we collected soil at three depths (rhizosphere, 5 cm, and 15 cm) associated with the invasive plant, *Psidium guajava* (guava), and native plant types. We determined that the sampling location (microclimate) was the strongest driver of both bacterial and fungal communities (74 and 38%, respectively), with additional minor but significant impacts from plant type and soil depth. This study highlights the continued need to explore microbial communities across diverse environments and demonstrates the weight of different abiotic and biotic factors impacting soil microbial communities across San Cristóbal Island in the Galápagos archipelago.

**IMPORTANCE/SIGNIFICANCE:** Human activity such as climate change, pollution, introduction of invasive species, and deforestation, poses a huge threat to biodiverse environments. Soil microbiota are an essential component to maintaining healthy ecosystems. However, a greater understanding of factors that alter these microbial communities is needed in order to find ways to mitigate and reverse the impacts imposed by human activity. The Galápagos Islands are a unique real-world laboratory, in that the islands’ biogeography and physical locations in the Pacific Ocean provide distinct microclimates within small geographic distances. Harnessing these distinct environments allowed us to investigate the influence of microclimates, soil depth, and vegetation cover on bacterial and fungal community composition.

## INTRODUCTION

Biodiversity is essential for sustaining proper ecosystem function^1^. However, due to global climate change, human impact (deforestation, introduction of invasive species, mining, population growth), and other factors, biodiversity across a vast array of environments is decreasing at an alarming rate^2^. Invasive species are one of the largest human-induced sources of global environmental change^3,4^. Much of the habitable world suffers from the introduction of invasive species; however, there are key locations that are particularly vulnerable^5,6^. The Pacific Islands, including the Galápagos Islands, are under intense threat from invasive plant species^7^. Roughly 1,400 invasive species have become naturalized in the Galápagos Islands, with the majority of them being plants or plant-associated organisms^8^.

The Galápagos Islands are a chain of islands of volcanic origin located ~1000 km west of the coast of Ecuador. One of the oldest islands in the archipelago is San Cristóbal. Due to their location along the equatorial Pacific, the islands experience distinct weather changes throughout the year, with major influences coming from the El Niño-Southern Oscillation^9^ and the Intertropical Convection Zone (ITCZ)^10^. The ITCZ largely dictates the seasonal weather patterns observed on the islands: January - June is the hot/wet season with large rain storms occurring across the entire islands while July - December is the cooler/dry season, with coastal zones experiencing rainfall and cloudy conditions at elevations around 700 meters above sea level (masl)^11,12^. This seasonal shift results in unique microclimates that are strongly dependent on elevation, ranging from arid near the coast to “transition zone” elevations to humid at the volcano summits^12^. These unique microclimates create distinct ecosystems over small geographic scales, which has led to speciation and the endemism of much of the Islands’ native flora and fauna. The high levels of endemism make the Galápagos Islands particularly susceptible to invasive species, where perturbations can quickly alter resident biodiversity and harm fragile ecosystems^13^.

Environmental microbes in soil and surrounding plants have the potential to greatly alter ecosystem functions^14,15^; however, microbes are often overlooked when discussing how to preserve biodiversity. Microbes play a pivotal role in global carbon and nitrogen cycling^16^, as well as in maintaining plant and soil health^17,18^ through the prevention of plant pathogens and pests^19^ as well as the secretion of metabolites that enhance plant growth^20,21^. Soil perturbations, such as those induced in agriculture and crop production, have been shown to reduce soil microbial diversity^22–24^. Although there have been extensive investigations into the factors that drive microbial communities, such as pH^25–27^, salinity^28–30^, soil depth^31^, soil organic carbon content^32^, temperature^33^, soil moisture^34^, redox status^35^ and plant communities^36–38^, these factors are likely environment specific; thus, continued investigation into the different drivers that apply to unique environments and ecosystems are still needed.

Unlike endemic flora, which are specialized for and adapted to specific microclimates, invasive flora are often able to tolerate a range of environmental conditions, providing them with a growth advantage that allows them to spread quickly. Beyond outcompeting endemic flora, invasive plants can negatively impact the soil nutrient levels as well as inhibit carbon and nitrogen cycling ^39^. However, while much is known about the impacts invasive plant species have on above-ground flora and fauna, we are still limited in our understanding of the role invasive plant species have on below-ground communities such as resident soil microbiota; many of plant-microbe interactions are expected to be plant-species specific^40–44^. The invasive plant *Psidium guajava* (common name: guava), is one of the most detrimental invasive plants on the Galápagos Islands. Unlike the native plants on San Cristóbal, which typically thrive in only a single microclimate, *P. guajava* can be found in most areas of San Cristóbal as well as other inhabited islands, except in the most arid locations on the coasts^45^.

The Galápagos Islands are uniquely suited to investigate the interplay of multiple factors on the composition of soil microbial communities. The physical features of the island landscape allow the impact of microclimate and plant species on soil microbial communities to be examined across distinct environments within a small geographic location, removing confounding variables typically present when larger physical distances must be examined to access distinct microclimates. In this study, we investigated soil microbial community structure on San Cristóbal Island in the Galápagos archipelago across arid, transition zone, and humid microclimates as monitored at the end of the wet season. We characterized and compared both the bacterial and fungal communities associated with the rhizosphere (the region of soil surrounding plant roots) of invasive and native plant types from three distinct microclimates, designing a study to examine these environmental variables as well as sampling depth. Our goal was to better understand the biotic and abiotic factors impacting the composition of soil microbial communities and their potential function. Such an understanding may allow us to harness the biological potential of these microbes to mitigate invasive plant species or alter the impacts of human activity on microbial communities. We determined that the strongest driver influencing both bacterial and fungal communities in the unique landscape of San Cristóbal was microclimate, with soil depth and plant species both modestly but significantly influencing these communities.

## RESULTS

### Sampling sites on San Cristóbal Island

To determine the effects of distinct microclimates and invasive plant species on resident soil microbiota in unique geographic locations, this study was conducted on the Island of San Cristóbal in the Galápagos Archipelago. We selected three sampling sites that encompassed three distinct microclimates within four miles of each other. The sites were: Mirador (leeward side, 320 masl, arid), Cerro Alto (leeward side, 520 masl elevation, transition zone), and El Junco (windward side, 690 masl elevation, very humid) (Figure 1A and Supplemental Figure 1A). These sites were chosen not only because they embody distinct environmental microclimates that are geographically close to one another, but also because they house weather stations that have been continuously collecting environmental variables since 2015, including temperature (°C), % relative humidity, and average rainfall (Figure 1B). Historically, Mirador is the warmest site in both rainy (January - June) and dry (July - December) seasons compared to the other two sites (Figure 1B). Cerro Alto experiences moderate temperatures and moderate rainfall throughout the year, while El Junco has the lowest average temperatures, highest % humidity, and highest average rainfall (Figure 1B).

**Figure 1.**
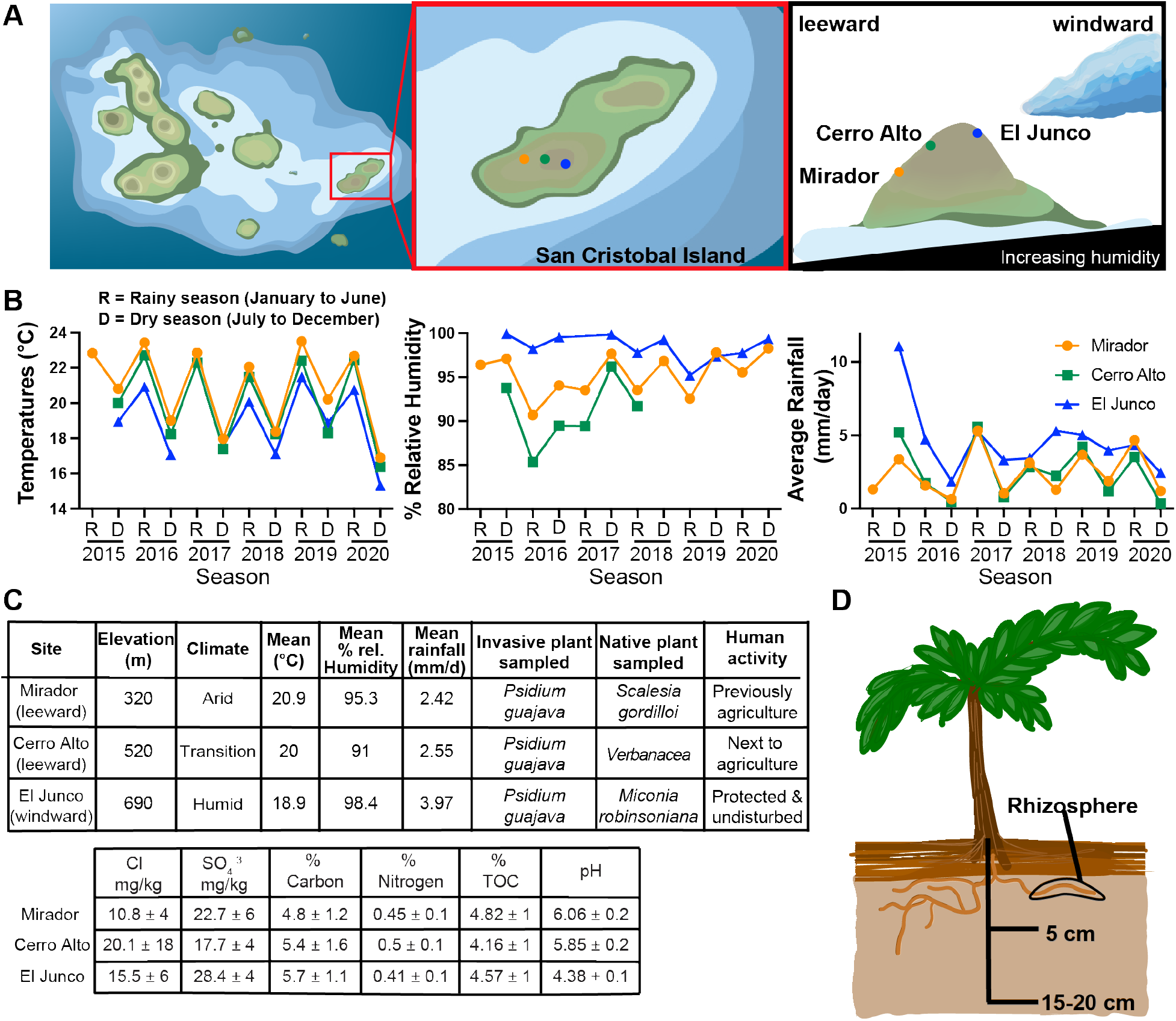
San Cristóbal Island in the Galápagos Archipelago exhibits geographically close sampling sites with distinct microclimates. (A) The relative locations of our three field sites on San Cristóbal in the Galápagos. (B) Temperature, % relative humidity, and precipitation data from weather stations installed in 2015. Data are the average of all collected over the given season. Due to technical issues arising from their remote locations, some weather station data is absent. (C) Tables detailing the microclimates (top) and soil properties (bottom) at Mirador, Cerro Alto, and El Junco. (D) Schematic of soil sampling scheme: rhizosphere, 5 cm, or 15 – 20 cm below the plant.

In addition to the three field sites having distinct microclimates, the vegetation, soil properties, and plants also varied with each site (Fig. 1). Prior to 2015, the land surrounding Mirador was used for agricultural purposes; it is now used as a conservation farm to grow native plants. The plot of land sampled at Mirador had several citrus trees and shrub grasses present with larger bushes, guava, and scalasia spaced out across the plot (Supplemental Figure 1A). The Cerro Alto site was densely packed with a range of plants including tall grasses, guava, and herbs; this land has not previously been used for agriculture, but it is near land with cow pastures. The land surrounding El Junco is protected by the Galápagos National Park; we sampled a plot containing guava, miconia, ferns, and small grasses (Supplemental Figure 1A).

### Soil sampling scheme

We performed soil collections at the end of the wet season, in June of 2019. At the Mirador and Cerro Alto sites, we obtained soil samples from six native and six non-native plants, while at El Junco we sampled from twelve of each plant type. The non-native plant, *Psidium guajava* (guava), was present at all sites. However, there was not a single native plant species sufficiently abundant to consistently sample at all three sites. Therefore, for each sampling site we selected a distinct native plant species that was similar in size to guava, abundant at that site, and within the same 50 m x 50 m sampling plot as the guava (Figure 1C): *Scalesia gordilloi* at Mirador, *Verbena* spp. at Cerro Alto, and *Miconia robinsoniana* at El Junco (Figure 1D and Supplemental Figure 1A). From each plant, we collected soil at three depths: (a) rhizosphere, the soil particles adhered to the plant root and shaken off (Supplemental Figure 1B), (b) 5 cm below surface from where the plant emerged from the soil, and (c) 15 to 20 cm below the plant (hereafter simply as 15 cm) (Figure 1D). The soil collected at each site differed visually in color, water content, and texture (Supplemental Figure 1C); we did not observe visual differences between different soil depths within a single sample site. We therefore collected 144 soil samples (Mirador and Cerro Alto each having six native and six non-native plants sampled at three depths and El Junco having 12 native and 12 non-native plants sampled at three depths).

### Soil carbon and nitrogen are influenced by plant type and soil depth

To better quantify the physiochemical differences between the soils at each site, we conducted soil analyses on pooled samples. From each site we pooled soils from the same plant type and sample depth (i.e., all guava from 5 cm; all miconia from rhizosphere, etc.) which led to 18 samples for soil analysis. When the data were analyzed by averaging all of the results from samples from the same site, we did not observe significant differences for %TOC (total organic carbon), %C, %N, or Cl^-^ mg/kg, but we did observe significant differences in SO4^2-^ mg/kg and pH (Figure 2A, Supplemental Table 1). El Junco had significantly elevated levels of sulfate and lower soil pH than the other sites (Figure 2A). When we then compared samples collected from guava versus the native plants across all sites, we observed significantly (*P* <0.05) elevated %TOC, %C, and %N in samples from guava compared to the native plant samples (Figure 2B). Additionally, we observed significantly lower %TOC, %N, and %C for the 15 cm samples compared to either the rhizosphere or 5 cm depth samples (Figure 2C). These data indicate that the invasive plant guava may either directly or indirectly impact soil carbon levels compared to the native flora. Additionally, soil closest to the plant, regardless of plant type, had higher levels of %TOC, N, and C compared to the 15 cm samples.

**Figure 2.**
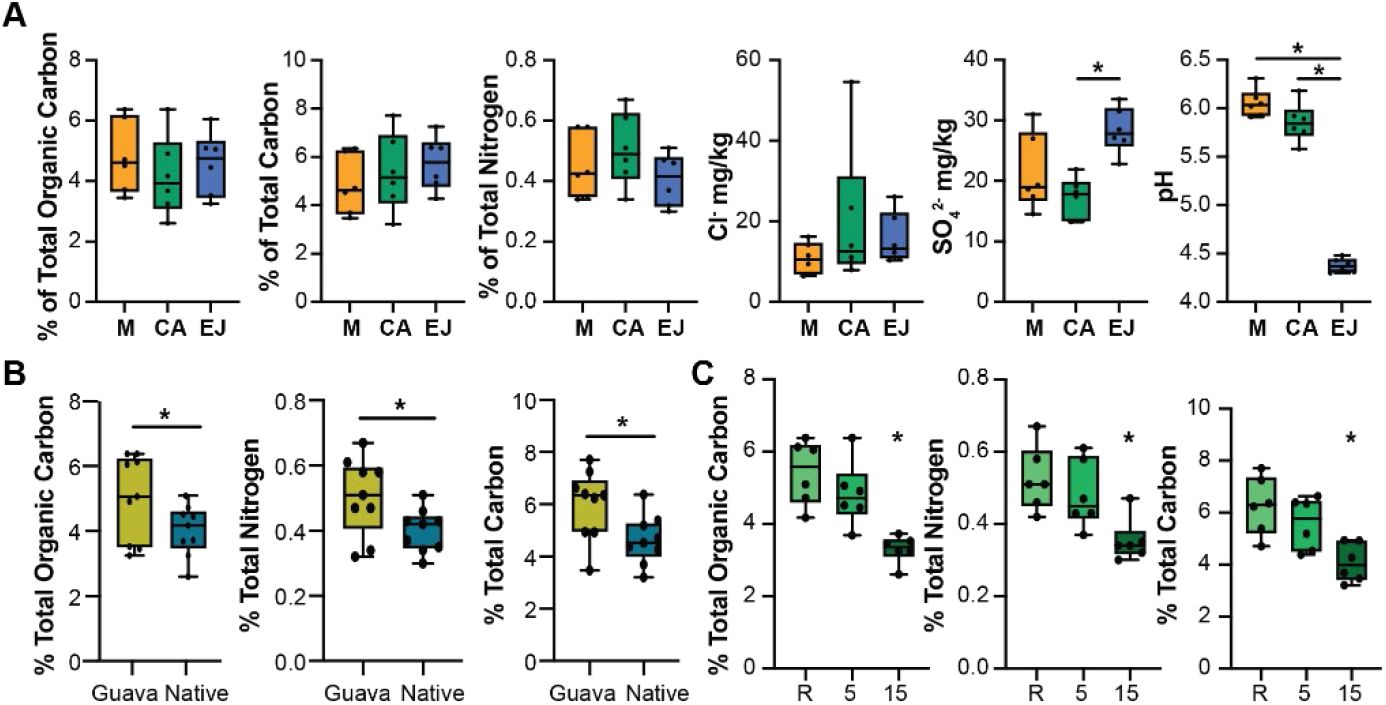
Soil properties vary by site, plant type, and soil depth. Samples from within each site, plant, and depth were pooled, resulting in 18 samples. A) Comparison of soil by site, averaged across other variables. B) Comparison of soil by plant type (guava vs. native; all native samples were averaged). C) Comparison of soil by depth (all samples were averaged by depth regardless of plant type or site). Each dot represents a independent sample. * denotes *P* <0.05; * above a group indicates that it is significantly different from all others on the graph. M = Mirador, CA = Cerro Alto, EJ = El Junco. R = rhizosphere, 5 = 5 cm depth, and 15 = 15 cm.

### The dominant bacterial and fungal taxa show distinct distribution patterns across sites

From the soil samples collected, we identified the soil microbial communities present using 16S rDNA and ITS (internal transcribed spacer) amplicon sequencing. Rarefaction analysis indicated sufficient read depth was achieved during sequencing (Supplementary Figure 2). We assigned bacterial taxonomy to amplicon sequence variants (ASVs) using a Naïve-Bayes classifier compared against a SILVA reference database that was trained on the V3-V4 region of the 16S rRNA gene^46^. When we assessed the relative abundances of bacteria and fungi at the phylum, order, and genus levels, we saw distinct distribution patterns of community composition at each site (Supplemental Figure 3). Furthermore, the overall patterns of both bacterial and fungal relative abundances were similar across all samples from a particular site, regardless of plant type or soil depth from which they were collected, indicating that each site has distinct microbial communities associated with it.

Across all three sites, the three dominant bacterial phyla were Proteobacteria (26%), Acidobacteriota (also known as Acidobacteria; 25%), and Actinobacteriota (also known as Actinobacteria; 20%) (Figure 2A). The most abundant phyla showed distinct distribution patterns across sites. The relative abundances of the Actinobacteriota, Methylomirabilota, Myxococcota, and Firmicutes were all highest at Mirador, intermediate at Cerro Alto, and then lowest at El Junco (Figure 3A). The Acidobacteriota and Proteobacteria exhibited the opposite pattern, with lowest relative abundances in the Mirador samples and highest in those from El Junco (Figure 3A). In contrast, the relative abundances of the Verrucomicrobiota and Bacteroidota were significantly higher at Cerro Alto than at Mirador or El Junco, while the relative abundances of the Plantcomycetota and Chloroflexi were lowest at Cerro Alto compared to both Mirador and El Junco (Figure 3A).

**Figure 3:**
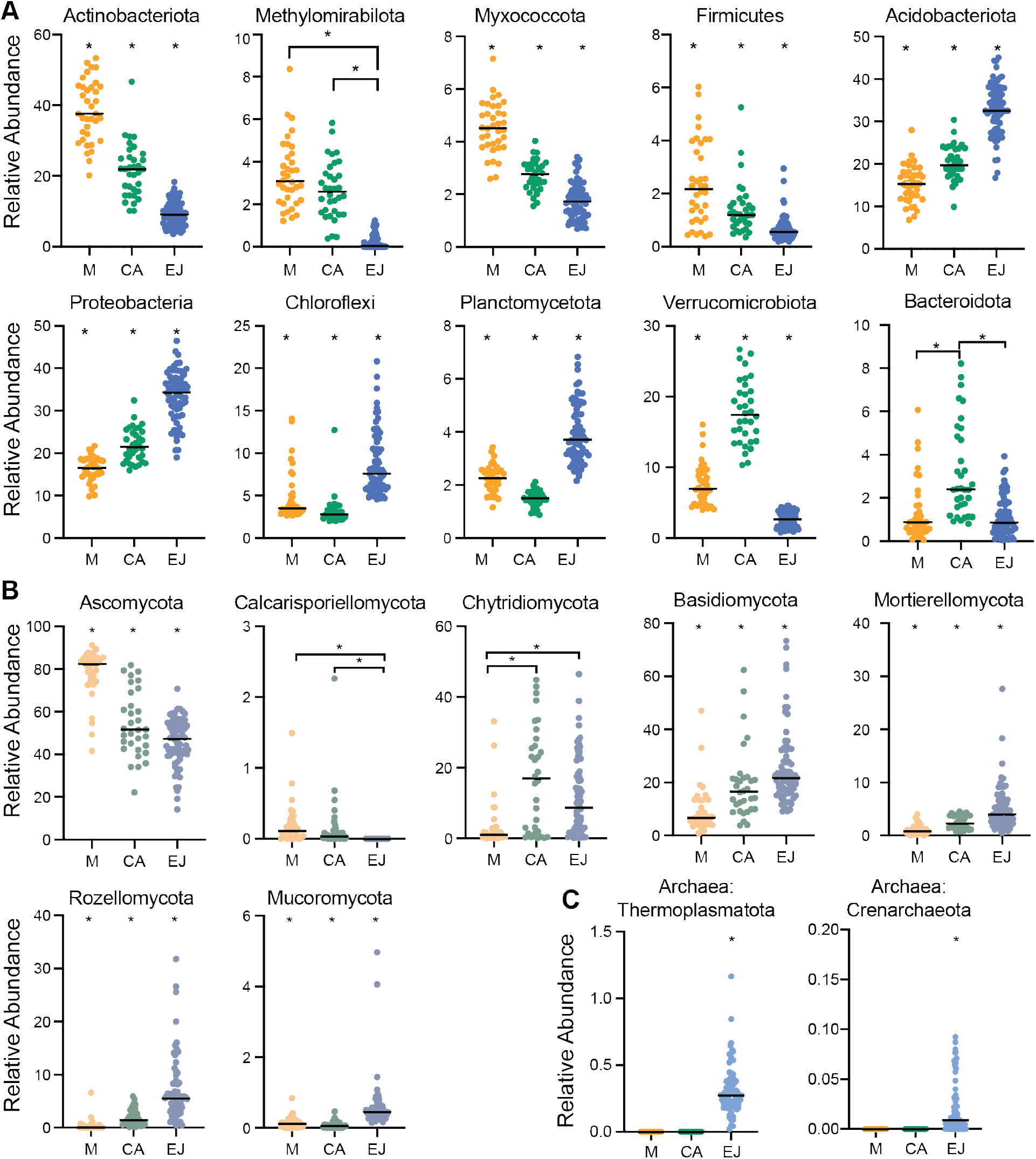
Dominant taxa have distinct distribution patterns. Relative abundances of (A) the eight most abundant bacterial phyla, (B) the seven most abundant fungal phyla, and C) two archaeal phyla from all samples from Mirador, Cerro Alto, and El Junco. Kruskal-Wallis ANOVA was used for initial statistical comparisons across all sites; once significance was identified, Mann-Whitney t-tests were used to compare between two sites. Asterisk (*) denotes significance *P* <0.05, if the asterisk does not have lines denoting the groups being compared, the group with the * is significantly different from all other groups in the graph.

We similarly assessed the relative abundance of the dominant fungal groups across the three sampling sites. The Ascomycota (56%), Basidomycota (20%), and Chytridiomycota (11%) were the dominant fungal phyla at all three sites, with the relative abundance of the Ascomycota being significantly higher at Mirador (78%) compared to Cerro Alto (54%) and El Junco (46%) (Figure 3B). The Basidiomycota and Mortierellomycota exhibited the opposite pattern, with their relative abundances being highest at El Junco (Figure 3B). Finally, the relative abundance of the Chytridiomycota was highest in Cerro Alto compared to the other two sites (Figure 3B).

Although we were not targeting archaea, we did detect two phyla using 16S rRNA sequences, the Thermoplasmatota and Crenarchaeota at El Junco, with undetectable levels at either Mirador or Cerro Alto (Figure 3C).

### The microbial communities at El Junco are the least diverse of the three sites

We next investigated the impact sampling site and plant type had on alpha diversity, or mean species richness. The Shannon entropy was similar at Mirador and Cerro Alto for both bacterial and fungal communities (*P* = 0.23 and *P* = 0.72, respectively) but it was significantly lower for the El Junco samples (*P* < 0.001 for bacterial and *P* < 0.01 for fungal communities) (Figure 4A), indicating that El Junco has significantly lower microbial alpha diversity than either Mirador or Cerro Alto.

**Figure 4:**
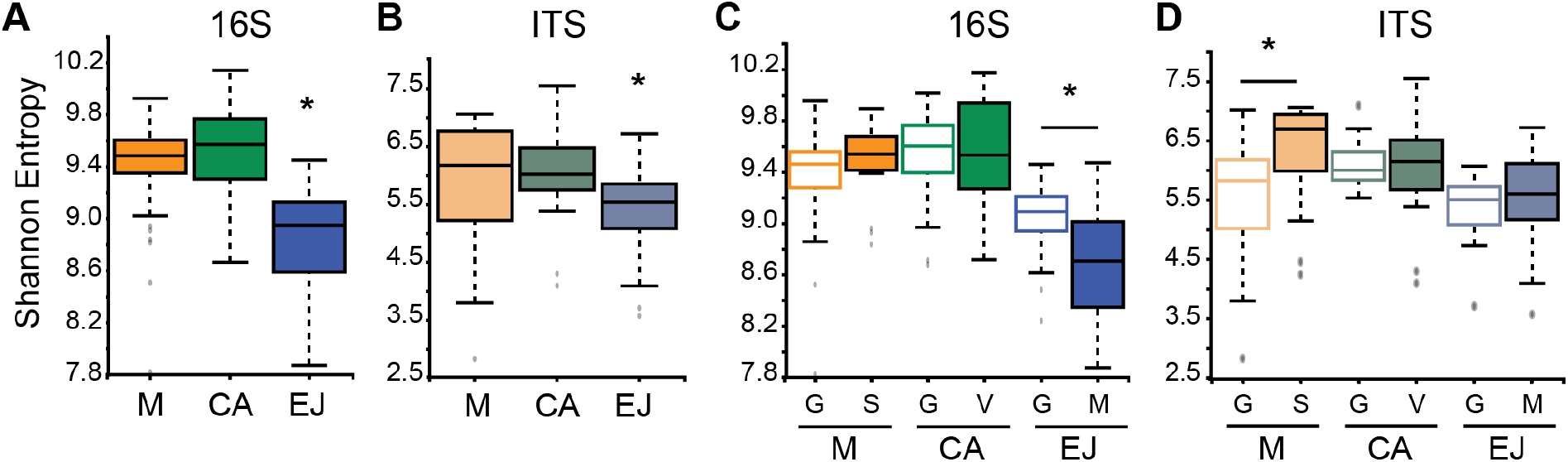
El Junco soil samples have a lower alpha diversity compared to samples from Mirador and Cerro Alto. Bacterial (A) and fungal (B) communities across sampling sites, and between the native and invasive plant samples within each individual site for bacterial (C) and fungal (D) communities. Alpha diversity was determined by measuring the Shannon Entropy for bacterial and fungal communities in all of our samples (Mirador n=36, Cerro Alto n=34, and El Junco n=72). Higher values indicate greater alpha diversity. Sites are coded by colors. Asterisk designates significance *P* < 0.01; if there is no bar between comparisons, this means that the group with the asterisk is significantly different than all others in that graph.

We then compared the alpha diversity of samples grouped by plant type (invasive or native) to investigate if the invasive guava had lower or higher diversity compared to the native plants at each site. At Mirador and Cerro Alto, there were no significant differences in bacterial alpha diversity between samples from each plant type (Figure 4C). However, at El Junco, guava-associated soil had significantly higher Shannon entropy (or bacterial diversity) than the native miconia-associated soil (*P* <0.001) (Figure 4A, right). In contrast, the Shannon entropy for the fungal communities at Cerro Alto and El Junco were similar, while at Mirador the native-associated soil had a significantly higher fungal diversity compared to guava-associated samples (*P* = 0.025) (Figure 4D). This indicates that, although microbial alpha diversity is not uniformly influenced by plant type, the bacterial communities at El Junco and the fungal communities at Mirador have distinct Shannon entropy values depending on the microbial association with the particular plant species.

### Geographic location predicts soil microbial community composition

To determine the major drivers of the microbial communities, we explored the variation in community composition across 142 samples (two Cerro Alto samples were excluded due to quality concerns). We visualized the relationships between the composition of all samples using non-metric multidimensional scaling (NMDS) plots based on the Bray-Curtis dissimilarity of the samples (Figures 5A and 5B). Samples in both the bacterial and fungal NMDS plots (Figure 5) clustered by geographic location, with ‘site’ explaining 74% of the observed bacterial variance and 38% of fungal variance (PERMANOVA, *P* < 0.001 for both). The microbial communities of El Junco soils were most distinct from those at Mirador and Cerro Alto, which had some minimal cluster overlap (Figure 5A and 5B). We observed significantly different bacterial and fungal beta diversity at each site (PERMANOVA, *P* = 0.001; Figure 5C) and an increasing taxonomic divergence between Mirador and the other two sites, Cerro Alto, and El Junco (PERMANOVA, *P* = 0.001) (Figure 5C), with the most dissimilar community from Mirador occurring at El Junco. Altogether, these data indicate that the composition of both the bacterial and fungal communities strongly correlates with site.

**Figure 5.**
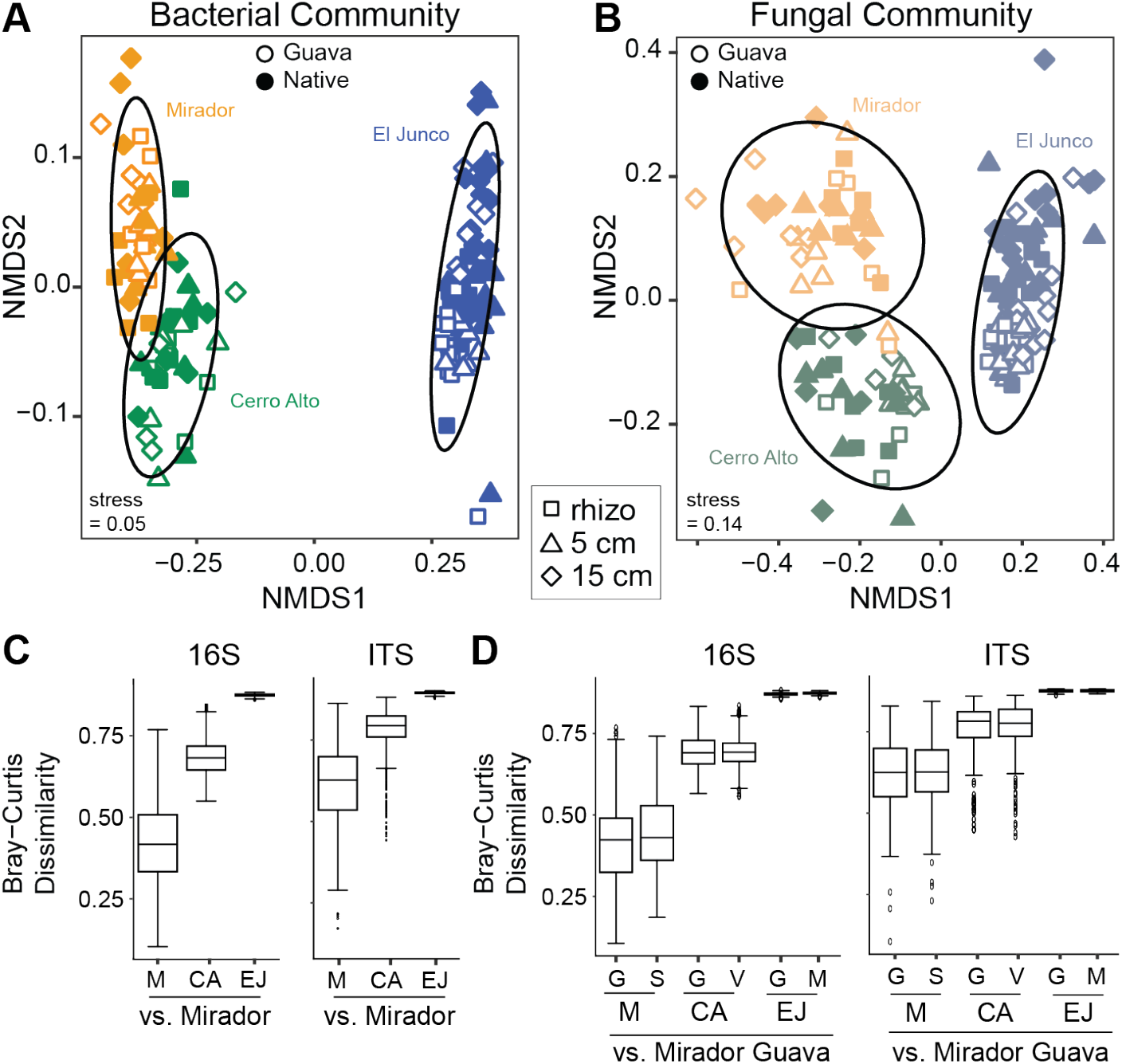
All three sites have significantly different beta diversity. We compared all (A) bacterial and (B) fungal communities. Ellipses indicate 95% confidence intervals. All sites are significantly different from each other (*P* < 0.05) by PERMANOVA. Beta diversity of (C) bacterial and fungal communities across microclimates, and (D) between the native plant and guava within each individual microclimate for bacterial and fungal communities. Beta diversity was determined by measuring the Bray-Curtis dissimilarity for bacterial and fungal communities in all of our samples (Mirador n = 36, Cerro Alto n = 34, and El Junco n = 72) and comparing the obtained values to Mirador. Higher values indicate greater beta diversity. The Bray-Curtis value of 0 indicates that the two communities are identical and share all taxa, and the value of 1 indicates that the two communities do not share any taxa.

### Soil depth and plant type also influence the soil microbiota

We then assessed the difference in beta diversity between samples associated with either the native or invasive (guava) plants and sampling depth at each site. We did not observe a significant impact of plant type on either the bacterial or fungal beta diversity (Figure 5D). However, the variance of the communities was significantly impacted by plant type, ranging from 8 - 16% (based on PERMANOVA analysis), with El Junco having the greatest plant-type-induced variance for both bacterial and fungal communities (Supplemental Figure 4). Additionally, comparing soil depth within each site indicated that soil depth did not influence Cerro Alto’s bacterial or fungal communities, but did impact Mirador and El Junco’s bacterial and fungal communities, explaining 7.8 - 31% of their variance (Supplemental Figure 5). The rhizosphere and 5 cm samples were more similar to each other than either was to the 15 cm samples (Supplemental Figure 5). Overall, these data indicate that while the bacterial and fungal communities are primarily influenced by geographic location, soil depth and plant species also affect their composition, with the extent of that effect dependent on sampling site.

### Dominant phyla are distinctly distributed between native and non-native sites

Having observed the strongest plant-induced variance at El Junco, we next wanted to compare the most abundant microbial phyla between guava and miconia samples at El Junco. We examined the ten most abundant bacterial and six most abundant fungal phyla in the rhizosphere and 5 cm soil samples (since these depths are most impacted by plant type, Supplemental Figure 5). The majority of the bacterial (8 out of 10) and fungal phyla (5 out of 6) relative abundances were significantly (*P* < 0.05) different between guava and miconia as determined by Mann-Whitney t-tests (Supplemental Figure 6A and 6B).

The phylum Glomeromycota was notable based on previous work on another Galápagos Island showing that invasive plants have a higher abundance of these fungi compared to native flora^47 48^. To determine if this trend was consistent across all sites, we compared the relative abundance of Glomeromycota from the rhizosphere and 5 cm-associated samples of guava and native plants at Mirador and Cerro Alto. However, there were no significant differences between the relative abundances of Glomeromycota based on plant type at either of these two sites (Supplemental Figure 6C). The relative abundance of Glomeromycota was similar in guava-associated samples across all sites (Supplemental Figure 6C), but a principal component analysis of the Glomeromycota ASVs showed that different genera of these fungi are present at each site (Supplemental Figure 6D)

### Predicted functional and ecological roles of the bacterial and fungal communities across the different sites

A major caveat to studying microbial communities is that differences in community composition may not lead to environmentally relevant functional differences. To gain insight into the functional and ecological potential of these soil communities, we used the predictors FARPROTAX for bacteria^49^, and FUNguild for fungi^50^. These predictors found that the dominant bacterial function were nitrogen fixation, nitrate reduction, chemoheterotrophy, and photoheterotrophy (Figure 6A and Supplemental Figure 7A) while the dominant fungal ecological guilds were saprotrophs, epiphytes, endophytes, and mycorrhizae (Figure 6B and Supplemental Figure 7B). These assigned categories were predominantly driven by sam pling site. Relative to the other two sites, El Junco was predicted to have the highest abundance of nitrogen fixers, with lower levels of nitrate reducers or denitrifiers, whereas Mirador was predicted to be dominated by nitrate reducers; Cerro Alto was predicted to have the highest abundance of denitrifiers, with moderate levels of nitrogen fixers or nitrate reducers (Figure 6A). In terms of carbohydrate degradation, each site was predicted to have distinct abundances of xylanolysis-active bacteria, while El Junco was distinct in its abundance of bacteria capable of cellulolysis, chitinolysis, and fermentation (Figure 6A).

**Figure 6:**
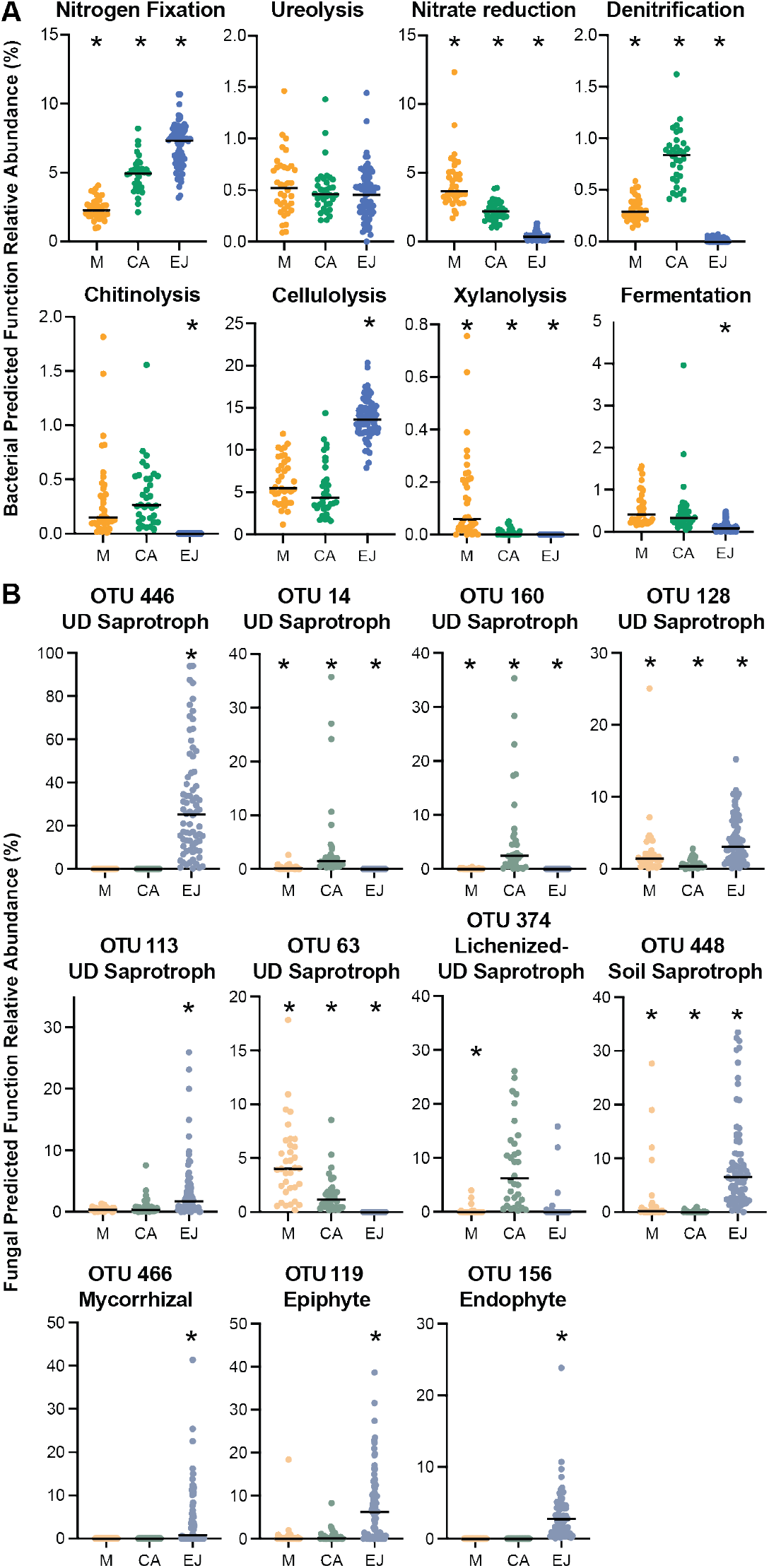
Functional and ecological potential of microbial communities across sampling sites. (A) Relative abundances of predicted functional potential of the bacterial communities and (B) fungal guilds communities at Mirador (M), Cerro Alto (CA), and El Junco (EJ). * = *P* < 0.05 for sites significantly different from the other two sites unless connected with lines. Significance was first determined by comparing all sites using Kruskal-Wallis ANOVA, and then pairwise comparisons using Mann-Whitney t-test.

For the fungal communities, the FUNguild predictions identified a number of prevalent saprotrophic operational taxonomic units (OTUs) that were distinctly abundant at multiple sites (Figure 6B). Multiple of these saprophyte OTUs (OTUs 446, 128, 113, 448) as well as other fungal guilds (mycorrhizal, epiphytic, and endophytic) were significantly enriched at El Junco relative to the other two sites (Figure 6B). In contrast, when we assessed the fungi predicted to be present at Mirador and Cerro Alto, they were instead dominated by fungi that fell into animal and plant pathogen guilds (Supplemental Figure 7B). Overall, these data suggest that not only are the microbial communities compositionally distinct at these three sample sites, but also that their predicted functional and ecological capabilities are distinct as well.

### El Junco guava-associated soil has an increased prevalence of free-living nitrogen fixers compared to miconia-associated soil

Based on El Junco having the highest predicted abundance of microbes involved in nitrogen fixation (Figure 6), we elected to further probe the microbial composition of the samples at this time. Specifically, since the guava-associated soil was found to have a high percentage of total nitrogen (Figure 2), we delved into the relative abundances of genera suggested to be free-living nitrogen fixers, including: Rhizobiums, Rhodomicrobiums, Roseiarcus, MND1, and Frankia. All of the genera investigated, with the exception of Rhodomicrobium, had significantly elevated relative abundances in guava-associated soil compared to miconia-associated soil at El Junco (*P* < 0.05) (Figure 7).

**Figure 7.**
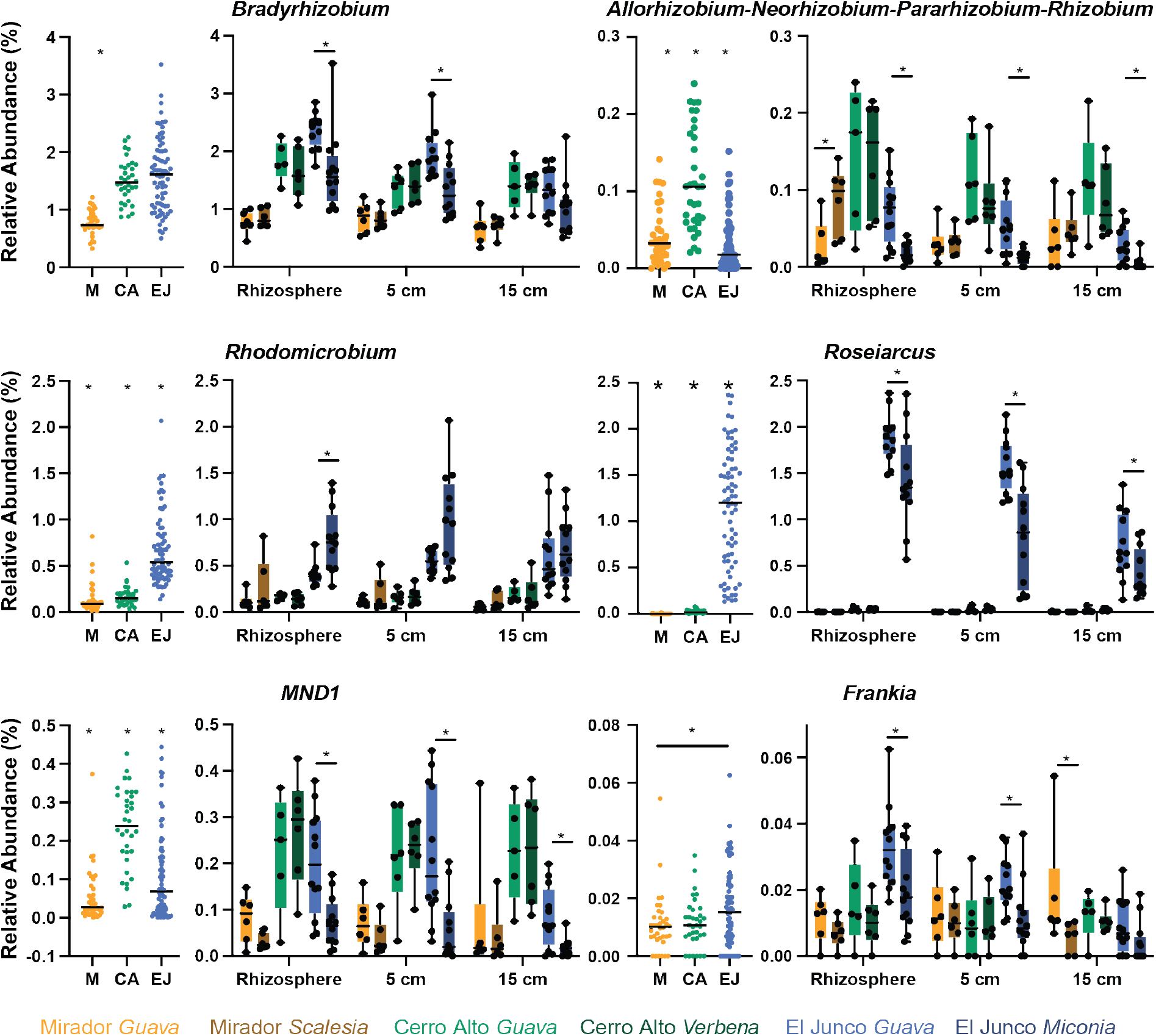
Distinct associations of six nitrogen fixing genera: *Bradyrhizobium, Allorhizobium-Neorhizobium-Pararhizobium-Rhizobium, Rhodomicrobium, Roseiarcus (Bekjerinckiaceae* family)*, MND1 (Nitrosomonadaceae* family), and *Frankia.* For each genus, the graph on the left indicates the relative abundances from all samples from Mirador, Cerro Alto, and El Junco; the graph on the right shows relative abundances of the genus for each sampling depth and plant type. Kruskal-Wallis ANOVA was used for initial statistical comparisons across sites. Mann-Whitney t-tests were sued to compare between two sites. Asterix (*) denotes significance P<0.05, the line denotes two groups being compared.

While computational predictors such as those based on 16S rRNA gene and ITS used above are suitable proxies for obtaining insights into the function of microbial communities, metagenomes, or all the genes present in a sample, are capable of providing deeper insights into the functions of the microorganisms associated with each sample. To further explore the functional component of the microbial communities associated with El Junco’s native and invasive plants, we therefore sequenced metagenomes from three randomly selected miconia and three randomly-selected guava-associated rhizosphere samples. Taxonomically, there were no differences detected between these six randomly selected samples, regardless of whether taxa identifications were derived from the 16S or ITS amplicon data or from the metagenomic data. We also did not see any statistical differences in canonical nitrogen-fixation genes. However, when looking at the predicted functions from the metagenomic samples, the samples associated with guava were distinct from those associated with miconia (Supplementary Figure 8).

## DISCUSSION

Elevation gradients are often exploited to investigate the environmental impacts on biodiversity. Previous work characterizing microbial community diversity has demonstrated conflicting impacts of elevation; some studies found higher microbial diversity at higher elevations^51,52^, while others found microbial diversity did not change with elevation^53,54^. These discrepancies in how elevation impacts soil microbiota may reflect the different environmental factors found at the specific sites and elevations being investigated^55^. Some of the most analogous studies to ours have been conducted in Hawaii. Both the Galápagos and Hawaiian Islands are volcanic in origin, forming as a result of hotspots beneath their respective tectonic plates. Although both of the above-sea level islands of these archipelagos are similar in age^56,57^, the soils at our sites are ~800,000 - 1,000,000 years in age^58^, while one Hawaiian study examining a temperature gradient was examining very young soils (~20,000 years old)^59^, and another examining an extensive elevation-precipitation gradient examined soils ~150,000 years in age^52^. In addition, the Galápagos tends to be cooler than expected for its equatorial location and the climate in the Galápagos is drier than in Hawaii, with fewer dust depositions from Asia^57^. Notably, our elevation resolution is sparse (since we only sampled from three elevations) compared to the larger number of samples taken in these Hawaiian studies^52,59^. Nevertheless, a comparison of our results is informative. One Hawaiian study examined sites where the mean annual temperature (as well as the carbon flux, but not the total carbon storage) varied, but where vegetation, soil type, moisture, and pH were constant^59^. No differences in bacteria richness was observed at these sites, indicating that other factors (i.e. pH, plants, precipitation) appear more important than temperature in driving bacterial community composition^59^. In the study of elevation-precipitation gradients on Hawaii, they observed that fungal richness increased with increasing precipitation, while bacterial richness appeared unimodal^52^. This is in contrast to our results, where both bacterial and fungal diversity metrics were lowest at the highest elevation (El Junco) compared to lower elevations. The relationship between pH and microbial diversity was also different between our work and that in Hawaii: in our study, higher pH correlated with higher bacterial and fungal diversity, while in Hawaii, fungal diversity was decreased at higher pH^52^. Overall, as a review by Looby and Martin 2020 points out, differences in biodiversity across topographic gradients are likely due to variations in a range of abiotic factors such as precipitation, landscape, and temperature^55^.

Abiotic factors previously shown to influence microbial community diversity and composition include pH^27,60,61^, temperature, soil moisture^27^, and organic content^27,62,63^. Here we determined that, overall, the site from which a sample is obtained (Mirador, Cerro Alto, or El Junco) accounts for ~70% of the bacterial and ~40 % of the fungal variance observed. El Junco having the lower overall diversity is consistent with prior studies indicating that acidic soils lead to lower bacterial richness^27^. In addition, El Junco is significantly more humid than the other two sites, and samples from this site exhibited the most distinct communities compared to Cerro Alto (transition zone) and Mirador (arid). Previous work exploring the influence of precipitation on microbial communities has demonstrated inconsistent patterns. In some cases, more humid environments have increased biodiversity compared to more arid sites^64,65^, while other studies have seen no changes in biodiversity metrics across precipitation gradients^66,67^. Cerro Alto, the site with the highest bacterial and fungal diversity, receives a moderate and fluctuating amount of precipitation. Previous work exploring wet-dry cycles and transition zones has indicated that differing microbial communities are present depending on whether the location is in the wet or dry part of the cycle^68–70^; this could explain the enhanced biodiversity observed at Cerro Alto. One limitation of environmental DNA sequencing is that sampling captures all DNA present in the sample regardless of whether the cells are dead, dormant, or alive. Thus, future work specifically investigating which microbial members are metabolically active during different seasons and environmental conditions (for instance with BONCAT or stable isotope probing^71,72^ along with manipulative experiments (such as altering water availability in soil cores) could provide additional insights into the influence of moisture and precipitation on microbial communities at these sites.

In addition to site-specific differences, we identified plant type as a minor but significant additional driver influencing microbial community variance between samples. Bacterial community composition varied between plant types in samples from Mirador (arid) and El Junco (humid), while the fungal community was influenced by plant type at all three sites. Plant-microbe interactions are a significant contributor to ecosystem functions and processes^73^, and the microbes involved in these interactions have the potential to enhance plant health^74–76^. However, how plant-microbe interactions shape the microbial community composition remains unclear, with some studies demonstrating plant type influences community^36–38,77^, while others do not observe a plant influence^78–81^, with one study showing that fungal endophytes varied over an elevation gradient, even from within a single plant host^82^. Plant roots excrete chemicals, nutrients, and carbon sources into the soil, which can alter microbial communities within the rhizosphere^83^. In our study, except for the bacterial communities at Cerro Alto, where the plant type does not significantly contribute to the observed community variance, we saw that approximately 9% of the bacterial and 12% of the fungal variance is explained by ‘plant type’, and that these values increase when selectively analyzing the samples closest to the plant root (rhizosphere and 5 cm).

Consistent with other soil microbiota studies^84,85^, the dominant bacterial phyla we observed were Acidobacteria, Actinobacteria, Bacteroidetes, Proteobacteria and Verrucomicrobia, and the abundance of these taxa tracked with different environmental variables present at each site. Actinobacteria, which form spores and thus are abundantly found in deserts^84,86^ were most abundant at Mirador, our most arid site. At El Junco, our most humid site, Proteobacteria and Acidobacteria were abundant, consistent with data showing they are less abundant in arid environments^84^. The acidic soil of El Junco (pH ~4.4) is consistent with the high prevalence of the acid-loving Acidobacteria found there^25,84,87,88^. In terms of fungi, the Ascomycota and Basidiomycota, frequently dominant phyla in soils^89–92^, were also abundant in our samples. Ascomycota are more abundant in more arid climates^89,93^, and were found at high levels at Mirador, while Basidiomycota are more abundant in high precipitation areas^52^ and were most abundant at El Junco. Thus, our data supports the idea that environmental factors such as soil moisture and pH are important determinants of microbial community composition.

Glomeromycota, which include the arbuscular mycorrhizal fungi (AMF)^94,95^, represented ~3% of the overall fungal abundance observed per site. AMF form an intricate symbiotic relationship with plants, enhancing the nutrient and water uptake ability of plants and their ability to tolerate stress, while the plant provides carbon to the AMF^96–101^. Approximately 90% of terrestrial plants worldwide form symbiotic relationships with AMF^102,103^. However, the native plants on the Galápagos Islands have significantly lower levels of AMF compared to similar plants on mainland Ecuador^48,104^ and invasive plants on Santa Cruz Island (another island in the Galápagos archipelago), had higher levels of AMF compared to native plants there; this was particularly true for guava^47,48^. Consistent with this, at the El Junco site, we found that soil associated with the invasive guava had significantly elevated levels of Glomeromycota compared to the native miconia; similar levels of Glomeromycota were observed between plants at Mirador and Cerro Alto. Although the addition of AMF is typically considered beneficial to the environment by restoring soil health and enhancing plant growth^105–107^, it is possible that in this environment, AMF could instead enhance the survival of invasive guava and thus be detrimental to native plants.

In addition to examining microbial diversity and abundance, it is possible to use amplicon data to gain insights into the potential functional roles of these communities using predictors such as FARPROTAX and FUNguild. While caveats of these approaches exist, in the absence of extensive (and expensive) SIP-metagenomic or meta-transcriptomic studies, these are reasonable analyses to obtain insights into these communities. Functional approaches also enable an examination of the potential functional redundancy existing within the population, a phenomenon where distinct species perform the same ecosystem function^108,109^; such functional redundancy is expected to increase with an increase in population biodiversity^110 111^. Our observation that plant and animal pathogens were predicted to be more abundant at Mirador and Cerro Alto relative to El Junco is consistent with Mirador previously being used for agricultural purposes and Cerro Alto being in close proximity to a current agricultural site, whereas El Junco is protected land distant from human activity. We also identified multiple congruent predictions at our most humid site, El Junco. Denitrification, nitrate reduction, and fermentation are all metabolic processes that occur in anaerobic conditions. All of these processes were predicted to be low at El Junco, indicating that these soils are likely largely aerobic. While our existing metagenomic data do not allow us to draw conclusions about the functional differences between sites, we expect that future work in this area will permit direct evaluation of microbial activity in these soils and thus a more nuanced understanding of their differences and similarities.

This study provides insight into the major predictors of soil microbial communities on the Island of San Cristóbal, Galápagos. We identified non-uniform correlations between elevation and environmental variables and microbial patterns at different geological and geographic locations. Because these volcanic islands and their soils are relatively young compared to continental soils, our findings suggest that microbial populations may consistently form under similar environmental conditions regardless of geographic location or relative age of landmass. The snapshot in time described here demonstrates that the bacterial and fungal soil communities at these sites are largely influenced by the microclimate and soil characteristics of each site, along with less dramatic effects by soil depth and plant type. The Galápagos Islands are expected to experience the impacts of climate change early relative to other areas, and thus are hot-spots to investigate this global environmental perturbation^112^. Future work investigating how these soil communities shift over time and in response to changes in the seasons is needed to address and potentially mitigate the effects of human-induced global changes such as climate change.

## ACKNOWLEDGEMENTS

This research was authorized under the agreement between the GSC-USFQ, Galápagos National Park and the Ministry of Environment of Ecuador with reference number MAE – DNB – CM – 2016 – 0041.

We would like to thank the UNC Center for Galápagos Studies (CGS) for funding provided to E.A.S. and D.R-I. through their Galápagos Seed Grant Program in 2018 and 2019 that allowed us to initiate and carry out these studies. We would also like to thank the administrators, staff, and scientists at the Galápagos Science Center for providing space, assistance, and equipment to carry out this work.

This material is based upon work supported by the U.S. Department of Energy, Office of Science, Office of Workforce Development for Teachers and Scientists, Office of Science Graduate Student Research (SCGSR) program (fellowship to S.M.Y). The SCGSR program is administered by the Oak Ridge Institute for Science and Education (ORISE) for the DOE. ORISE is managed by ORAU under contract number DE-SC0014664. All opinions expressed in this paper are the author’s and do not necessarily reflect the policies and views of DOE, ORAU, or ORISE.

A portion of this research was performed on a project award (10.46936/expl.proj.2019.51105/60000139, to E.A.S.) from the Environmental Molecular Sciences Laboratory, a DOE Office of Science User Facility sponsored by the Biological and Environmental Research program under Contract No. DE-AC05-76RL01830.

We would like to thank the Wellesley College Undergraduate Student Summer Research Program for providing funding to support C.M. work on this research.

## MATERIALS AND METHODS

### Soil sample collection

Soil samples were collected at three sites on San Cristóbal Island, Galápagos Islands, Ecuador on consecutive days in June 2019. The three collection sites were: Mirador (June 20, 2019; lat: - 0.886068, long: −89.539818), Cerro Alto (June 21, 2019; lat: −0.883333, long: −89.516667) and El Junco (June 22, 2019; lat: −0.8966, long: −89.480206). At Mirador, we collected soil surrounding 6 guava and 6 scalasia plants. At Cerro Alto, we collected soil surrounding 6 guava and 6 verbena, and at El Junco, we collected soil surrounding 12 guava and 12 miconia plants. Enhanced sampling was performed at El Junco as there were sufficient numbers of plant species within our 50 m x 50 m plot. Additionally, this site was the most preserved and had limited human impact thus, we wanted to have enough power to determine if invasive plants had unique or altered microbial communities compared to native plants. All equipment and gloves were wiped down between sampling using isopropanol wipes. From each plant we collected roughly 5 g of soil from three depths per plant (rhizosphere, 5 cm below ground, and 15 - 20 cm below ground surface). To collect soil from the rhizosphere, we pulled roots out and shook off soil into a 5 mL conical tube. To collect 5 cm and 15 - 20 cm sample depths, we measured the depth from the surface and collected plugs horizontally from the soil. Collections took less than 4 hours at each site and samples were stored in a cooler with ice packs for transport until returning to the lab, where samples were allotted for microbiological analysis and frozen at −20 °C until further processing.

### DNA isolation

After transport from the collection site on ice, soil samples for DNA isolation were aliquoted (~1 g into 1.5 ml Eppendorf tubes) and stored immediately at −20 °C. DNA isolation was performed using Qiagen DNeasy PowerSoil Kit (Cat # 12888-100), according to the manufacturer’s instructions with slight modifications: samples were vortexed for 30 min at maximum speed in the PowerBead tubes, and DNA was eluted in 50 μl of Solution C6. Isolated DNA yield and quality was checked using a Spectrophotometer – Thermo Scientific NanoDrop 2000 (Cat #ND-2000). DNA extracts were transferred on ice to the high-throughput sequencing facility at University of North Carolina at Chapel Hill for amplicon sequencing or to EMSL for metagenomic sequencing.

### Amplicon 16S and ITS rRNA gene sequencing

Members of bacterial and fungal communities were identified by sequencing the V3-V4 region of the 16S rRNA gene and the ITS region, respectively. Soil samples were sequenced according to the protocol described by Caporaso et al.^113^. The extracted DNA was amplified with barcoded primers to enable multiplexed sequencing. We used 341F 5’-CCTACGGGNGGCWGCAG-3’ and 806R 5’-GACTACHVGGGTWTCTAAT-3 for bacteria and ITS-9F 5’-GAACGCAGCRAAIIGYGA-3’ and ITS-4R 5’-TCCTCCGCTTATTGATATGC-3’ for fungi. The 16S and ITS libraries were constructed with the same barcodes for each sample and combined into the same amplicon pool after they were purified using AMPure beads (Beckman Coulter, Illinois, USA), pooled into a library (100 ng) and quantified by qPCR. Twenty percent of denatured PhiX was added to the amplicon pool (12 pM) and sequenced on the MiSeq platform using 300+300 bp paired-end V3 chemistry (Illumina, San Diego, CA). Raw sequence files were deposited to the NCBI Sequence Read Archive under the BioProject accession number: XXX.

### Metagenomic sequencing

The template library was generated using Nextera XT Library prep kit (FC-131-1024) according to the manufacture protocol. Pair-end sequencing with read length of 150bp was done on NextSeq550 platform using NextSeq 500/550 High Output v2 kit 300 cycles (cat#20024908). Shotgun metagenomic sequences were analyzed using the bioBakery suite of computational tools^114^. First, KneadData (v0.7.7) was used to perform quality control of raw sequence reads, such as read trimming and removal of reads matching a human genome reference. Next, MetaPhlAn (v3.0.7, using database mpa_v30_CHOCOPhlAn_201901) was used to generate taxonomic profiles by aligning reads to a reference database of marker genes. Finally, HUMAnN (v3.0.0a4 with databases) was used to functionally profile the metagenomes^115^. Sequences are deposited in the NCBI database XXX under the accession number XXX

### Processing of the sequence data

We processed sequences in QIIME 2 v2020.8^116^, using a modified pipeline workflow^117^. Primers were removed using the cutadapt plugin^118^ and the reads were sorted based on the primers into either the 16S (bacterial) or ITS (fungal) group. Raw sequence reads from soil samples were denoised, filtered and clustered into amplicon sequence variants (ASVs) using the Divisive Amplicon Denoising Algorithm (DADA2) plugin^119^. For the bacterial sequence data, we used the following parameters: ASVs with a frequency of <0.1% of the mean sample depth were removed to account for a bleed-through between MiSeq runs, and the remaining reads were used for subsequent analysis. For the 16S rRNA gene dataset, sequences belonging to mitochondria and chloroplasts were filtered out. After denoising and filtering, the number of sequences per sample ranged between 50 and 555,568 sequences for the 16S run and 7 to 105,710 sequences for the ITS run. Samples with fewer than 20,000 sequences were excluded from the analysis. These included samples M6-A, C3-C, and C1-B for the 16S rRNA gene amplicon dataset and C1-C and C6-A for the fungal dataset. A multiple-sequence alignment was created using MAFFT, and FastTree was used to create an unrooted phylogenetic tree, both with default values^120^. Taxonomy was assigned to the ASVs using a Naïve-Bayes classifier compared against SILVA v138.99 reference database trained on the 515-926 region of the 16S rRNA gene^46^. For fungal ASVs, Naïve-Bayes classifier was compared against the UNITE (ver8_99_s_04.02.2020) ITS database files.

To visualize the community composition, we constructed bubble plots and stacked bar plots of bacterial relative abundances using the ggplot2 package in R v4.0.3^121^. Alpha-diversity was estimated using Shannon diversity index^122^. Bray-Curtis dissimilarity was used to assess compositional differences between conditions (sites or plants or soil depth). The clustering of bacterial communities was visualized via NMDS plots constructed from Bray-Curtis distance matrices using ggplot2^121^. We used Permutational Multivariate Analysis of Variance (PERMANOVA) tests to determine whether the microclimate, sampling depth, or the plant species shaped microbial community structure^123^ using the adonis function from the R vegan package^124^. We used Bray-Curtis dissimilarity with 999 permutations. The strata term was used to limit permutations within individual sites to account for repeated measurements. We compared across both experiments, but we also ran PERMANOVAs separately for each of the microclimates.

### Functional group characterization and core taxa identification

To predict the fungal guild or functional groups, we used FunGUILD v1.0, an open annotation tool that predicts ecological guild by parsing fungal OTUs^50^. To visualize these results, we constructed a stacked bar plot of guild relative abundances using the ggplot2 package in R v4.0.3. To predict the functional potential of the soil microbiome from 16S rRNA genes, we used FAPROTAX^49^. FAPROTAX uses experimental evidence, such as culturing, to estimate metabolic phenotypes based on the 16S rRNA gene. To identify core taxa associated with the rhizosphere of each plant or site, we used COREMIC^125^.

#### Differential abundance

Associations of bacterial and fungal phyla across different sites or plants was evaluated by using DESeq2^126^ in MicrobiomeDB^127^. DESeq2 uses an empirical Bayesian approach to fit abundance data to a negative binomial model and converts P values for differential abundance to q values to correct for multiple hypothesis testing.

#### Weather stations

Weather stations were deployed in 2015 at each of the sampling sites (Images of weather stations from Mirador and Cerro Alto in Supplemental Figure 1). The weather stations collected data on precipitation, wind speed and wind direction, temperature, and solar radiation. Data gaps were the result of instrument failure. Due to remote location and difficult terrain, repairs to instrumentation took time.

#### Soil Analysis

Soil was collected and dried at 56 °C for 72 hours and then moved to room-temp and air-dried for an additional 2 weeks. Soil was then stored in plastic ziploc bags at room temperature in the dark until soil analysis was performed. We pooled ~1 g from each of the 144 by site, depth, and plant type (ex. El Junco guava rhizosphere samples) resulting in 18 samples total. Soil analysis was performed by the University of Georgia Athens, Agriculture, and Environmental Service Lab. The Dumas method was used for the analysis of total Carbon and Nitrogen and previously described^128^, briefly, for total organic carbon soil is treated with water and 8% sulfuric acid, mixed, and incubated for several hours and then placed in the soil dryer to volatilize carbonates from the sample. Soil (~0.5 g) is loaded into a steel crucible and combusted in an oxygen atmosphere at 1200°C in a Elementar Vario Max Total Combustion Analyzer. Elemental C and N are converted into CO_2_, NO_x_, and N_2_. Carbon content was determined using an infrared cell and N2 was determined using a thermal conductivity cell. Results were reported as a %. To measure anions USEPA Method 300 was used as previously described () briefly, a Dionex Model DX-120 ion chromatograph (Sunnyvale, CA, USA) was used with a 25 μl sample loop injector and AS40 automated sampler employed along with a Dionex IonPac AS14A separation column (3 × 150 mm) and IonPac AG14A guard column (4 × 50 mm). A Dionex ACRS-500 (2 mm) Anion Self-Regenerating Suppressor was used for conductivity suppression before detection by a Dionex DS4-1 conductivity cell. Soil was mixed at a 1:1 ratio in deionized water for an hour and filtered before running the samples. Eluent: 8 mM Na_2_CO_3_ + 1 mM NaHCO_3_ (1 ml min-1) was added at a flow rate of 0.48 ml per minute.

---

**Supplementary Figure 1:**
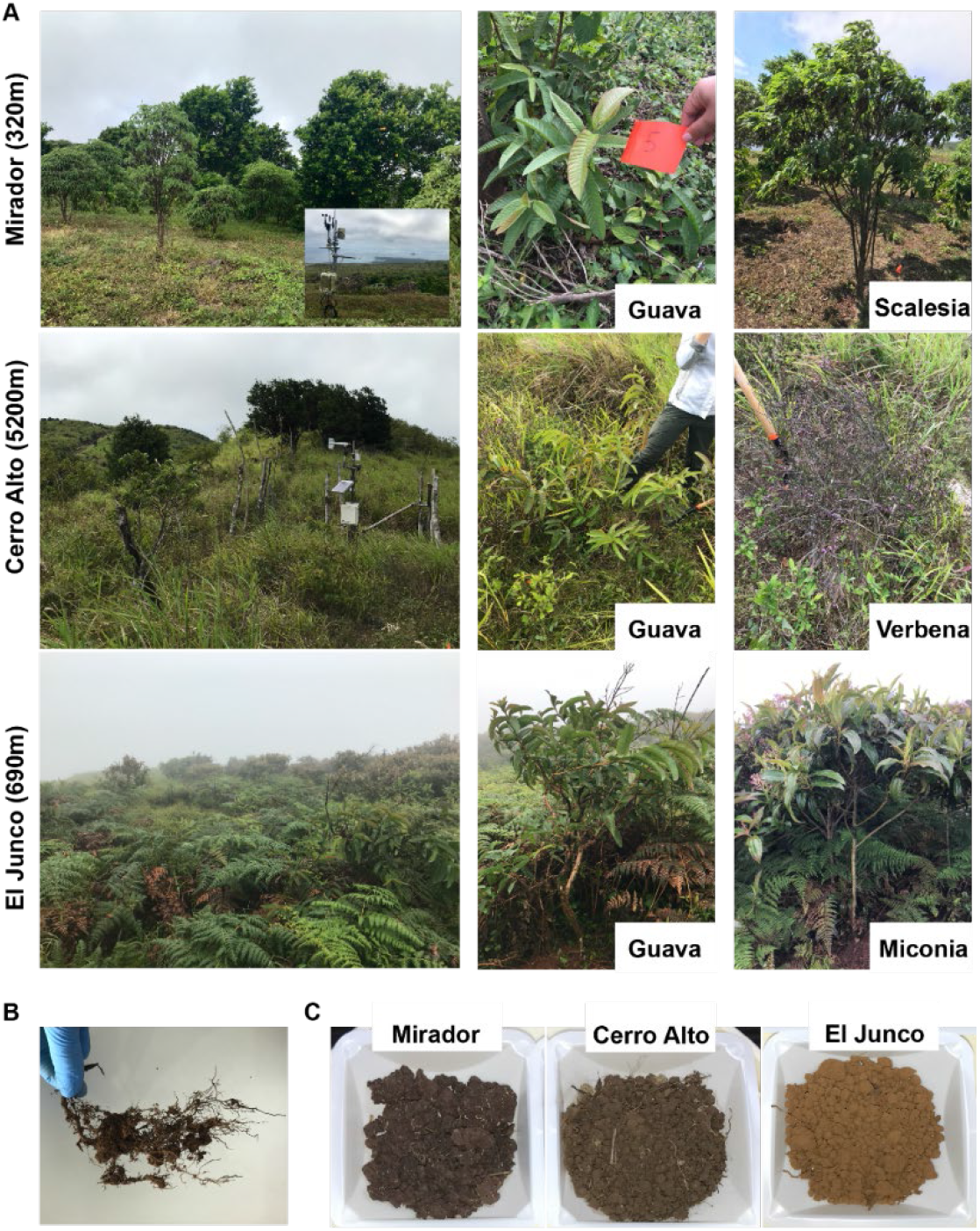
Images of field sites and plant species located at each site. (A) Field photos taken in June-July 2019 of sampling sites and plants sampled at each location; Mirador (guava and scalesia), Cerro Alto (guava and verbena), and El Junco (guava and miconia). (B) Image of root-associated soil. C) Images of the different soils from each sampling site.

**Supplementary Figure 2.**
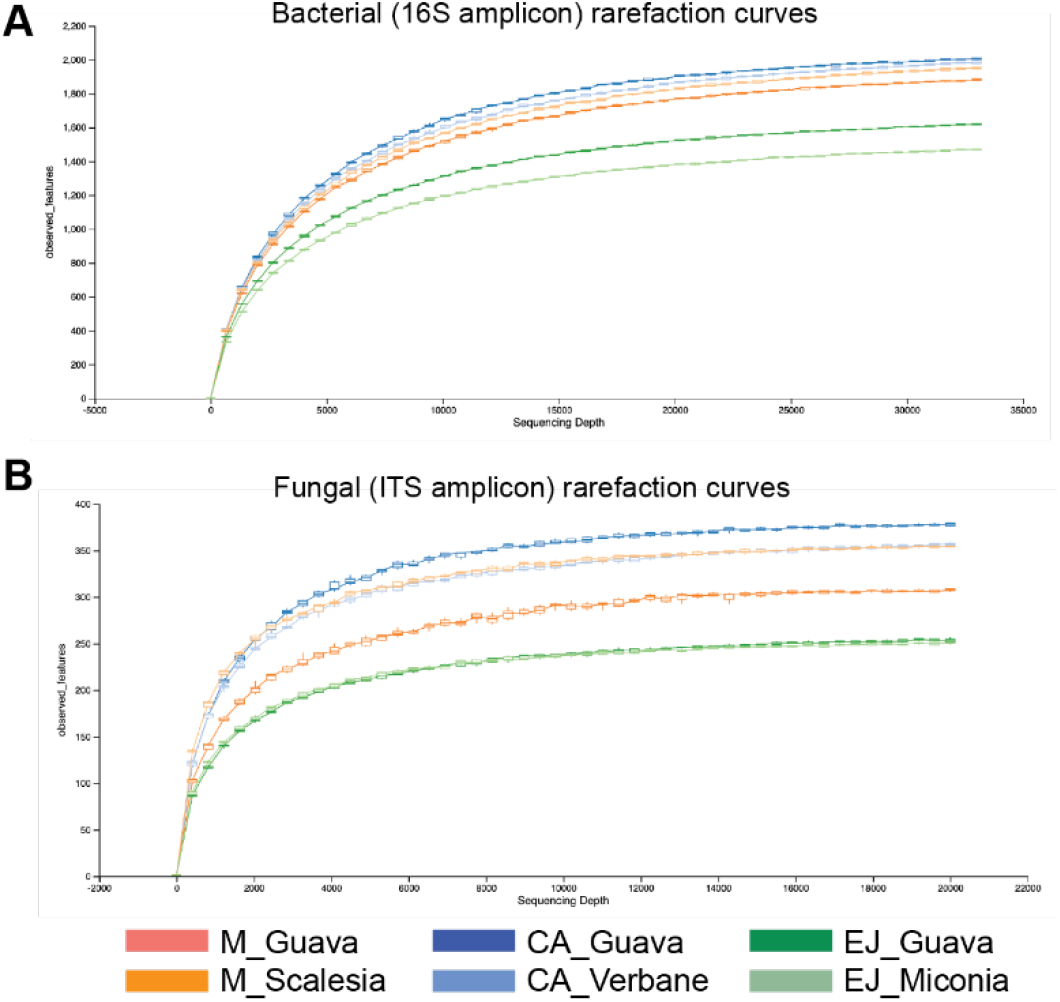
Rarefaction analysis of amplicon sequences. Rarefaction curves of A) Bacterial (16S) and B) Fungal (ITS) denoting analysis at each site and each plant were sufficiently sampled and reached asymptote.

**Supplementary Figure 3:**
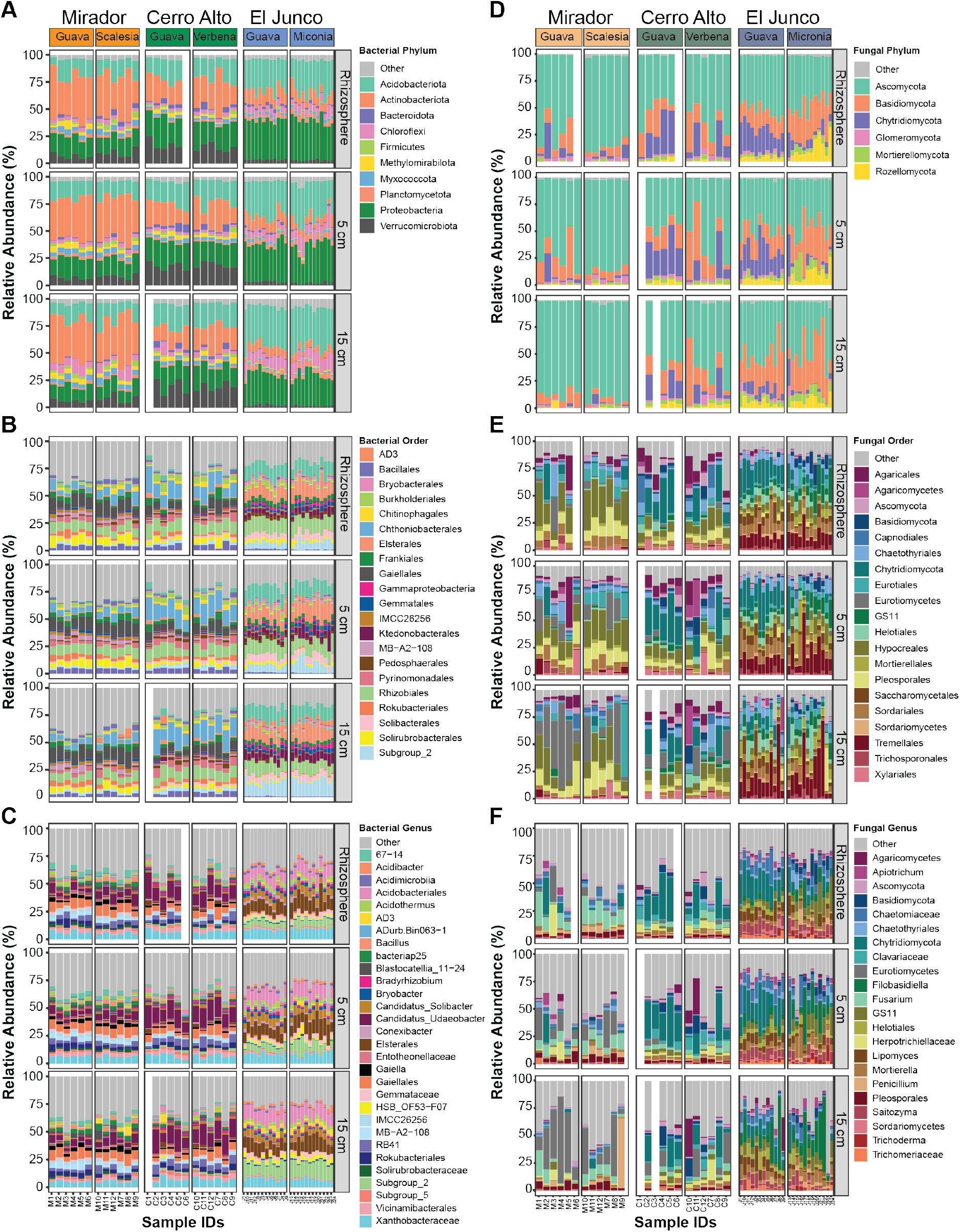
Relative abundances of bacterial and fungal communities at each sampled site. Samples are grouped by site, plant, and depth. Stacked bar plots show the relative frequency of 16S rRNA (left) and ITS (right) gene sequences assigned to each (A, D) phylum, (B, E) order, and (C, E) genus across samples. Each color represents a specific phylogenetic designation. Only taxa with relative abundances above 1% are shown; all others are grouped into the Other category. Blank bars represent missing data.

**Supplementary Figure 4.**
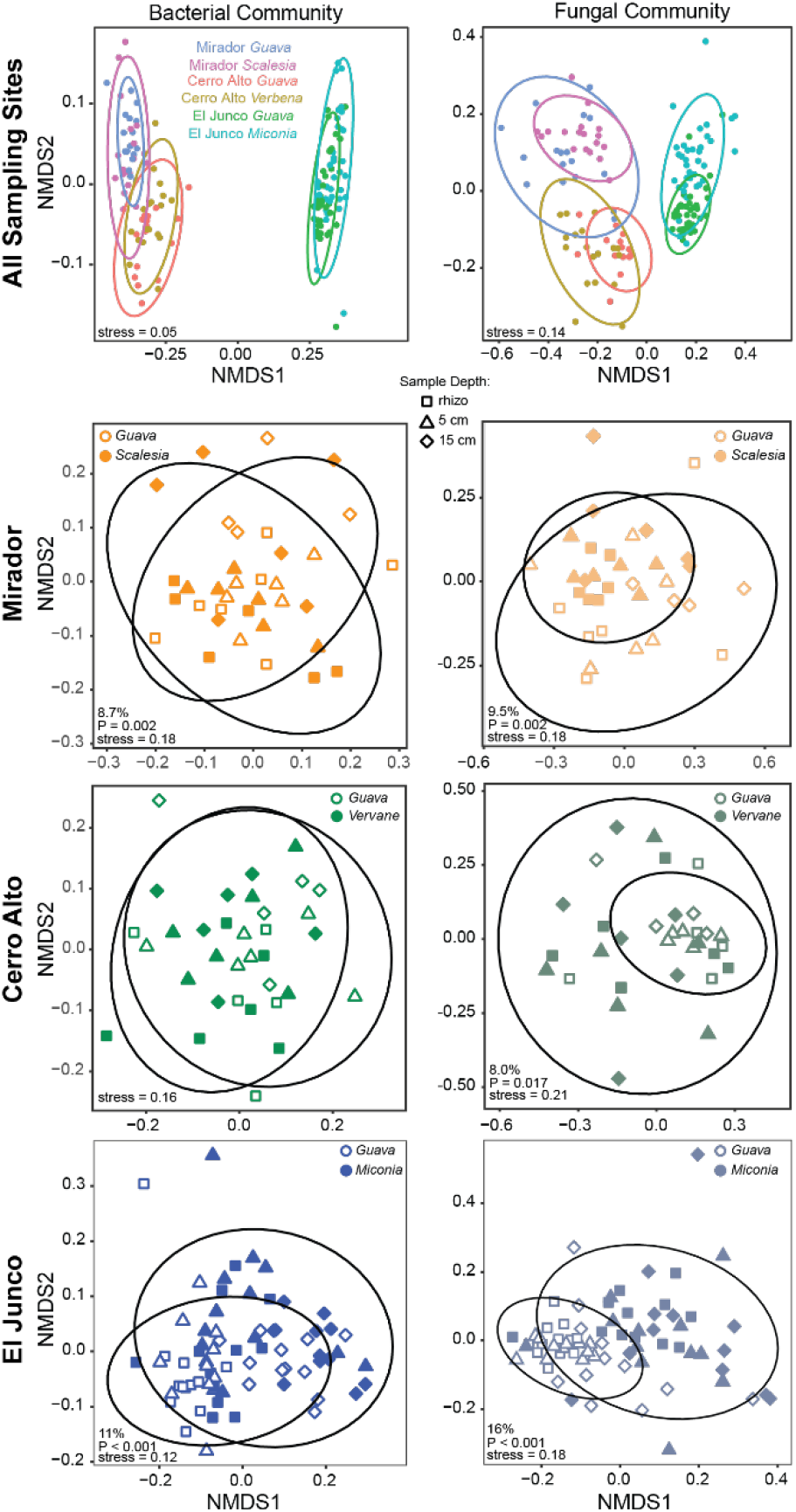
Plant species influence soil microbiota. NMDS plots of Bray-Curtis distances to compare all samples by plant type at all sites. Bacterial communities are shown on the left and fungal communities on the right. Ellipses indicate 95% confidence intervals for samples from each plant type. Community variance attributable to soil depth and its associated *P* and NMDS-stress values are shown in the bottom corner of each graph.

**Supplementary Figure 5:**
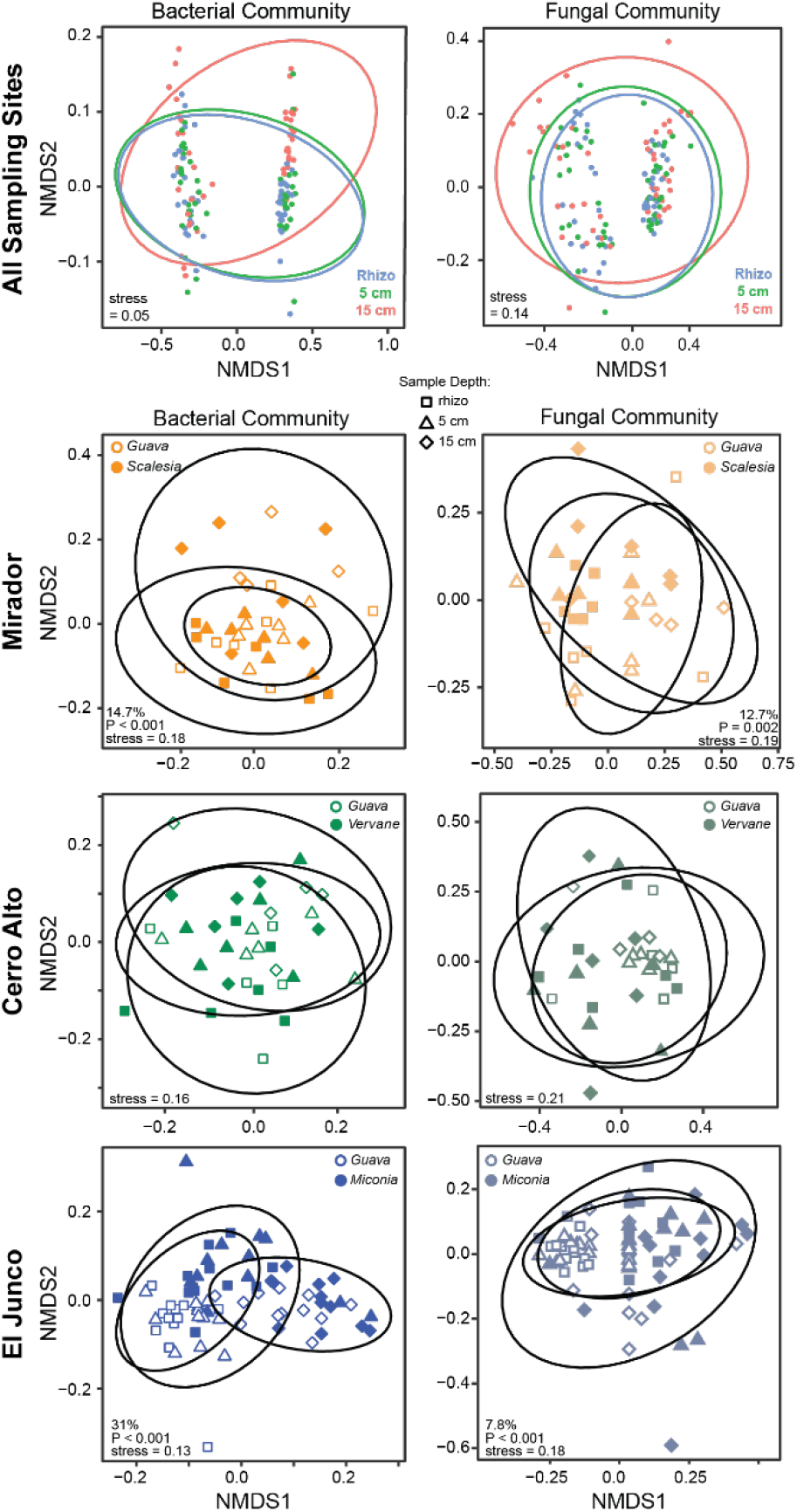
Soil depth influences soil microbiota. (Top) NMDS plots of Bray-Curtis distances comparing all samples by sample depths. Bacterial communities are shown on the left and fungal communities on the right. (Bottom) NMDS plots of Bray-Curtis distances comparing sample depths within each indicated site. Solid symbols = native plant; open symbols = guava. Ellipses indicate 95% confidence intervals and are grouped by depth. Community variance attributable to soil depth and its associated *P* and NMDS-stress values are shown in the bottom corner of each graph.

**Supplementary Figure 6:**
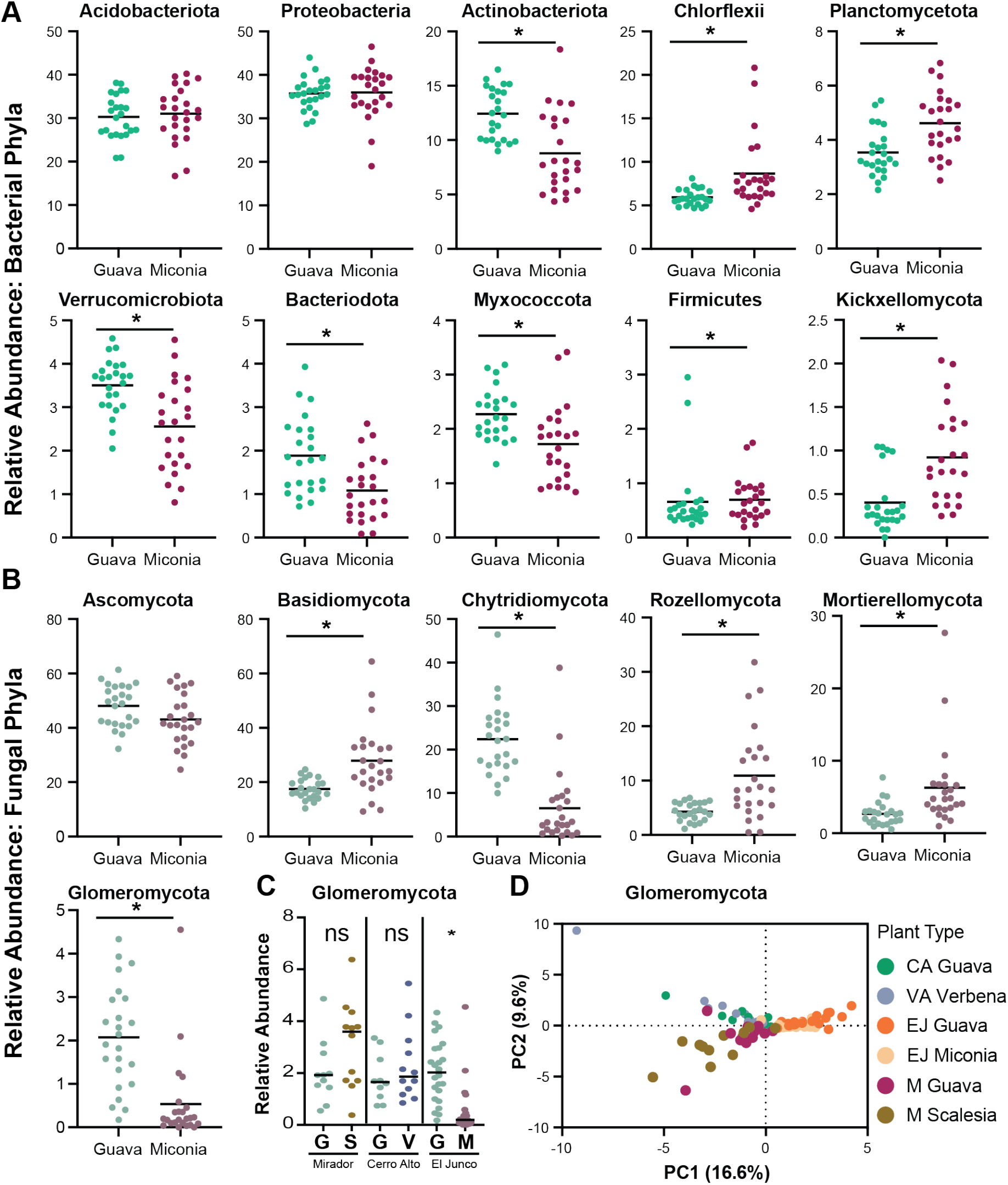
Relative abundances of bacterial and fungal phyla from guava and miconia at El Junco. (A) The ten most abundant bacterial and (B) six most abundant fungal phyla of samples from the rhizosphere and 5 cm soil associated with either guava or miconia at El Junco. (C) Relative abundance of Glomeromycota from the rhizosphere and 5 cm soil samples from either guava or native plant at Mirador, Cerro Alto, and El Junco. (D) Principal component analysis of ASVs in the Glomeromycota phylum. The relative abundance was compared using a Mann-Whitney test. Asterisk (*) denotes significance *P* <0.05.

**Supplementary Figure 7:**
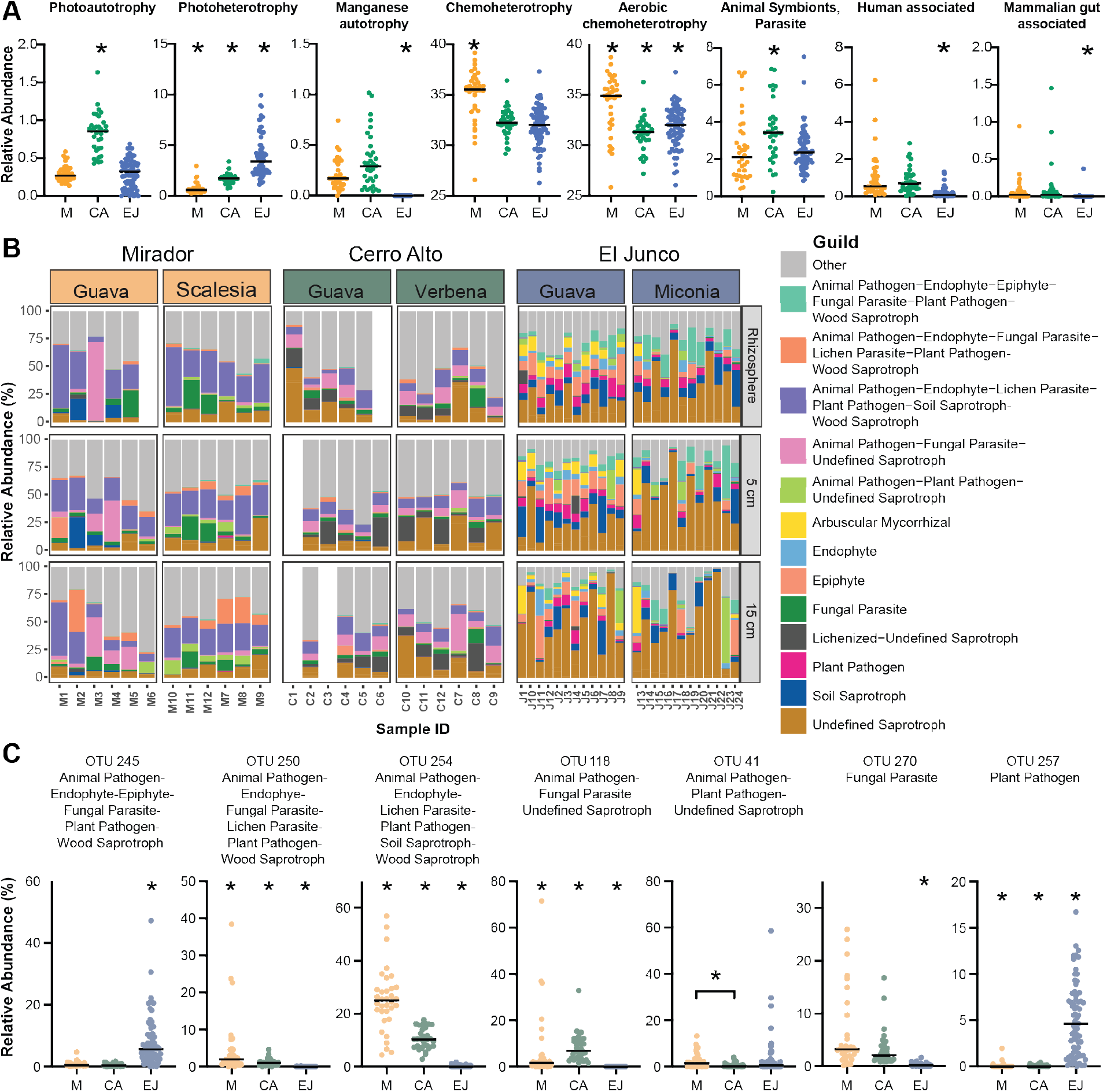
(A) Relative abundances of predicted functional potential of the bacterial communities at Mirador (M), Cerro Alto (CA), and El Junco (EJ). (B) Relative abundances and variation of fungal guilds across different sampling depths, plants, and sites based upon fungal *genus-level* classification. Each color represents a fungal guild. Samples are grouped by plant within each panel. (C) Relative abundances of fungal guilds at M, CA, and EJ. Asterisk (* = *P* < 0.05) denotes the site that is significantly different from the other two sites, unless two sites are connected with lines. Significance was determined by first comparing all sites using Kruskal-Wallis ANOVA, and then performing pairwise comparisons using Mann-Whitney t-test.

**Supplementary Figure 8.**
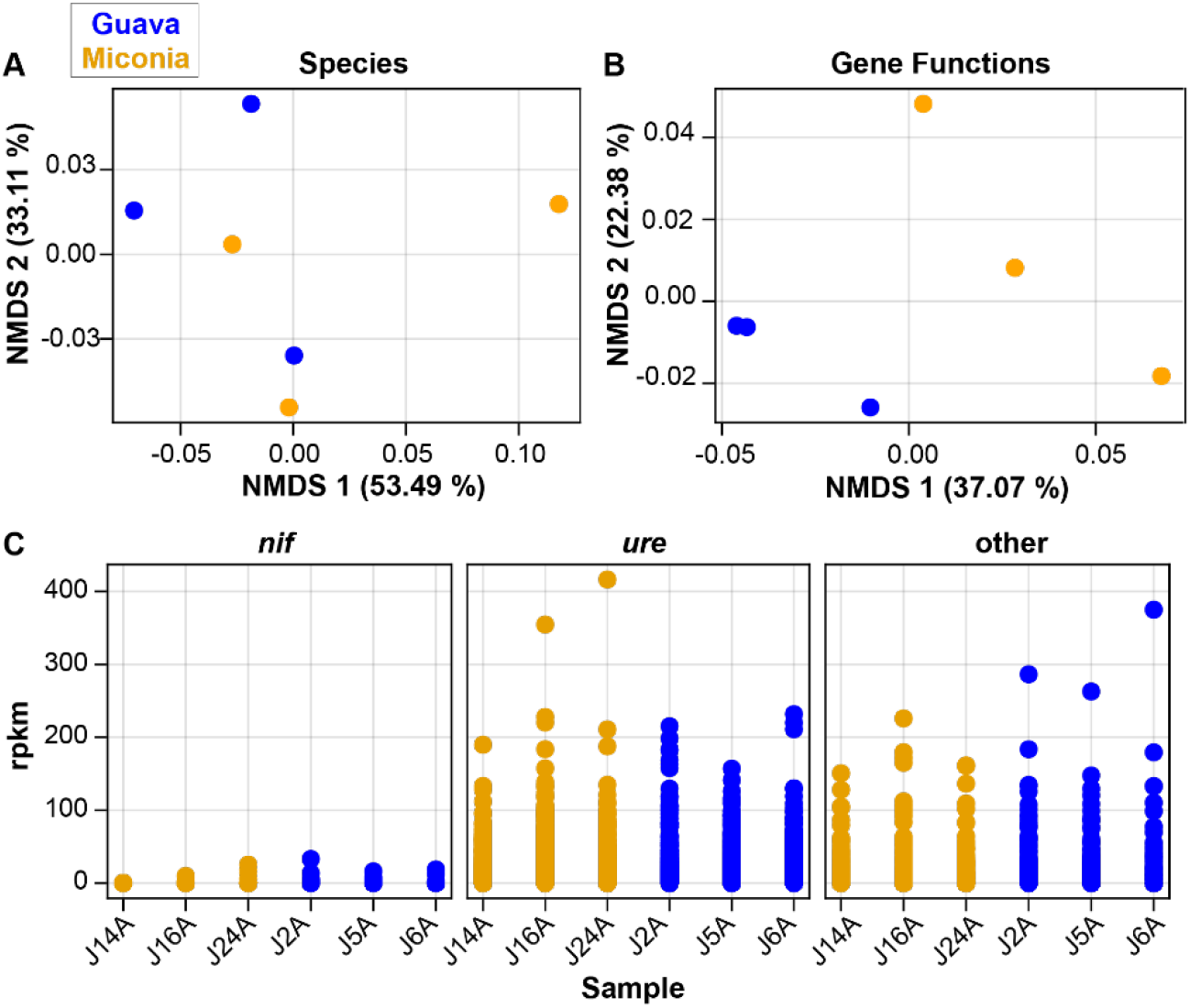
Taxonomic and functional profiling comparing native and invasive species rhizobiomes. Three samples each from guava and miconia were analyzed using shotgun metagenomic sequencing. A) Principal coordinate analysis (PCoA) using classical multidimensional scaling of Bray-Curtis dissimilarity in taxonomic profiles. Fourteen species in total were identified, 7 of which were found in all samples (mean prevalence = 64%). Three species (*Dyella* sp. DHG54, *Nitrolancea hollandica,* and *Limnochorda pilosa*) were found only on miconia, and 3 (*Actinospica robiniae, Bradyrhizobium erythrophlei,* Candidatus *Solibacter usitatus*) were found only on guava. B) PCoA of functional profiles, after removing reads that could not be assigned to any known gene function (mean 84.7%). Of 837,903 identified genes, 206,699 (24.7%) were found in all samples (mean prevalence 53.2%), 3,846 (0.46%) were found only in miconia, and 13,020 (1.55%) were found only in guava. C) Genes (matching reads per kilobase per million reads, RPKM) identified as *nif* (nitrogen fixation), *ure* (urease - nitrogen recycling), or other nitrogen reduction genes.

**Supplemental Table 1.**
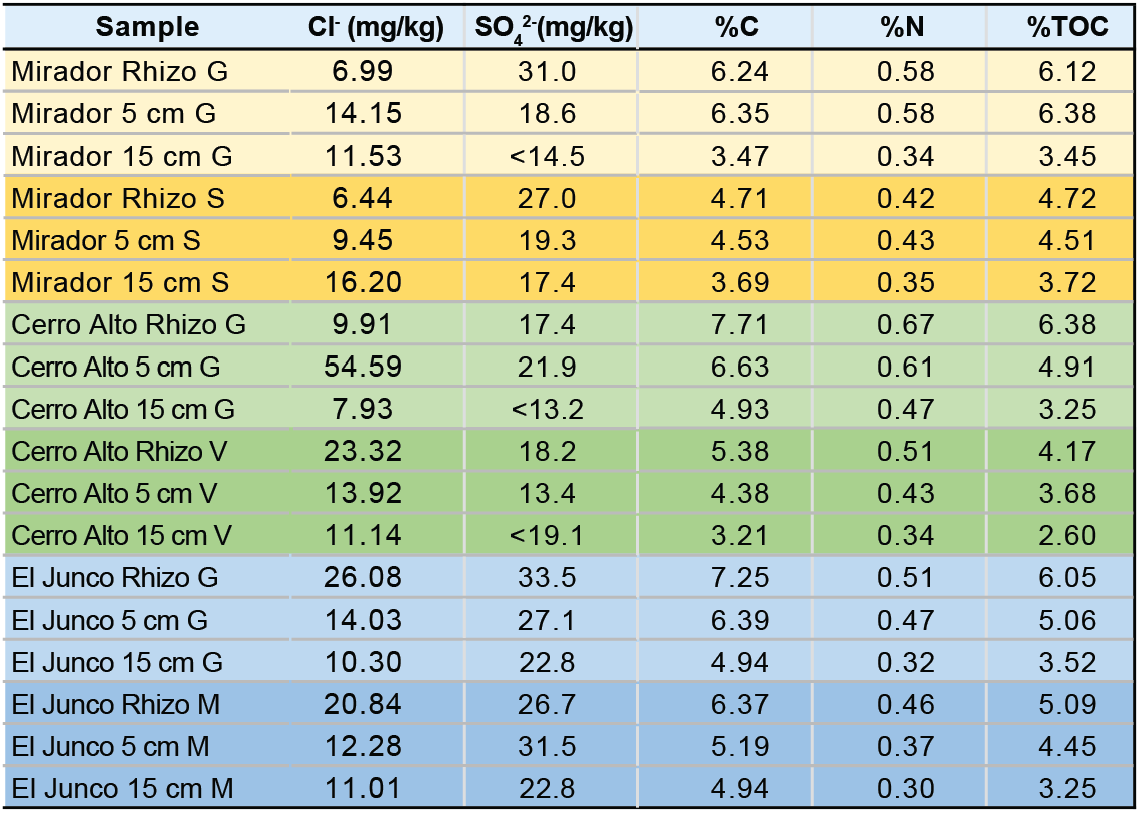
Soil nutrients above detectable limits. Samples from each site, plant, and depth were pooled, resulting in 18 samples total. Soil analyses were performed to measure Cl^-^, SO4^-2^, PO4^-3^, NO3N, F^-^, %TOC, %C, and %N. (PO4^-3^, NO3N, and F^-^ were measured but below detectable limits). Rhizo, 5 cm, and 15 cm refer to sample depth. Plant type is indicated by: G = guava, S = Scalesia, V = Verbena, and M = Miconia.

